# Contrasting patterns of specificity and transfer in human odor discrimination learning

**DOI:** 10.1101/2024.09.10.612215

**Authors:** Xiaoyue Chang, Huibang Tan, Jiehui Niu, Kaiqi Yuan, Rui Chen, Wen Zhou

## Abstract

Practice enhances olfactory performance. However, laboratory studies to date suggest that olfactory learning is largely restricted to the trained odors, posing a significant challenge for training-based rehabilitation therapies for olfactory loss. In this study, we introduce various types of odors to olfactory discrimination training, conducted unilaterally. We demonstrate contrasting patterns of specificity and transfer of learning, independent of adaptation and task difficulty. Individuals trained with odor mixtures of different ratios show long-term perceptual gains that completely transfer to the untrained nostril and effectively generalize to untrained mixtures dissimilar in structure and odor quality from the trained ones. Conversely, those trained with odor enantiomers show no transfer of learning across nostrils or to unrelated enantiomers, replicating our earlier findings (Feng & Zhou, 2019). Our observations indicate that concentration ratio and chirality represent distinct olfactory attributes. Furthermore, discrimination learning occurs at different stages of olfactory processing, depending on which attribute is task-relevant. These findings open up new avenues to enhance the effectiveness of olfactory training.

## Introduction

Olfactory training is recommended for patients with olfactory loss caused by infectious, traumatic, or neurodegenerative conditions (Hummel et al., 2017). It has received widespread attention due to the surge of patients with olfactory dysfunction from the COVID-19 pandemic (Tan et al., 2022). The typical routine of olfactory training in clinical research and practice involves twice-daily exposure to four single compounds –– phenylethyl alcohol, eucalyptol, citronellal, and eugenol –– with the odors of rose, eucalyptus, lemon, and clove, respectively, for 12 or more weeks (Hummel et al., 2009). However, the effectiveness of this protocol is not always obvious (Lechner et al., 2022) and is sometimes attributed to spontaneous recovery or the placebo effect (Yan, 2023). Little is known about the underlying mechanism for improvement following olfactory training. Generally, it is assumed that training fosters neural plasticity, particularly adult neurogenesis in the peripheral olfactory pathway (e.g., olfactory epithelium) (Hummel et al., 2017; Moreno et al., 2009; Wang et al., 2004).

A large body of literature on the phenomenology and mechanisms of visual perceptual learning (Fahle, 2005; Gilbert & Li, 2012; Lu & Dosher, 2022; Seitz & Dinse, 2007; Watanabe & Sasaki, 2015) indicates that it occurs at multiple levels and persists over weeks and months. The magnitude, specificity, and transfer of learning depend on factors such as stimuli, task, reinforcement, attention, and training protocol. The patterns of specificity inform about the locus of neural plasticity. This research has shed light on the brain processes underlying perceptual learning and laid the foundation for developing and optimizing rehabilitation therapies for amblyopia, myopia, and low vision (Chung, 2011; Deveau et al., 2013; Durrie & McMinn, 2007; Rodan et al., 2022). In contrast, only a few laboratory studies have examined the specificity/transfer of olfactory perceptual learning. They reported that repeatedly exposing one nostril to the steroid androstenone increased detection accuracy of androstenone in both nostrils for individuals who initially could not perceive its odor (Mainland et al., 2002). Repeated test exposures (threshold testing) to everyday odor compounds enhanced olfactory sensitivity only in women of reproductive age and in an odorant-specific manner (Dalton et al., 2002). Prolonged discrimination training (with feedback) in one nostril using a pair of odor enantiomers produced a gain in chiral discrimination that did not transfer to the other nostril or to structurally-dissimilar odor enantiomers (in terms of the chiral center) (Feng & Zhou, 2019). No formal assessment was conducted on how long-lasting the learning effects were. No generalization of learning to odorants unrelated to the specific training material was ever observed.

For olfactory training to be optimally useful, it should generalize to untrained odorants and provide long-term benefits. This means that learning needs to occur at a stage where odor representations are unaffected by the nostril of origin or differences in chemical structure or perceptual features between trained and untrained odorants. In other words, the training task and odorants need to engage high-level olfaction. These considerations led us to examine the specificity, transfer, and persistence of olfactory perceptual learning of odor mixture discrimination. We chose discrimination over passive odor exposure and odor mixtures over single compounds because complex olfactory tasks and stimuli are more likely to involve high-level olfactory processing and thus promote transfer of learning. Additionally, most real-world odors are mixtures that do not smell exactly alike. For comparison, we also conducted discrimination training with odor enantiomers and assessed the learning effect.

## Results

### Contrasting patterns of specificity and transfer in olfactory learning to discriminate mixtures and enantiomers

The experimental procedure in Experiment 1 included three phases (Figure 1A): baseline, unilateral olfactory training, and post-training testing. Participants, both men and women, were randomly allocated into two groups of 12 individuals each. Over a period of several weeks, one group was trained and tested with odor mixtures (mixture group), and the other with odor enantiomers (enantiomer group).

**Figure 1.**
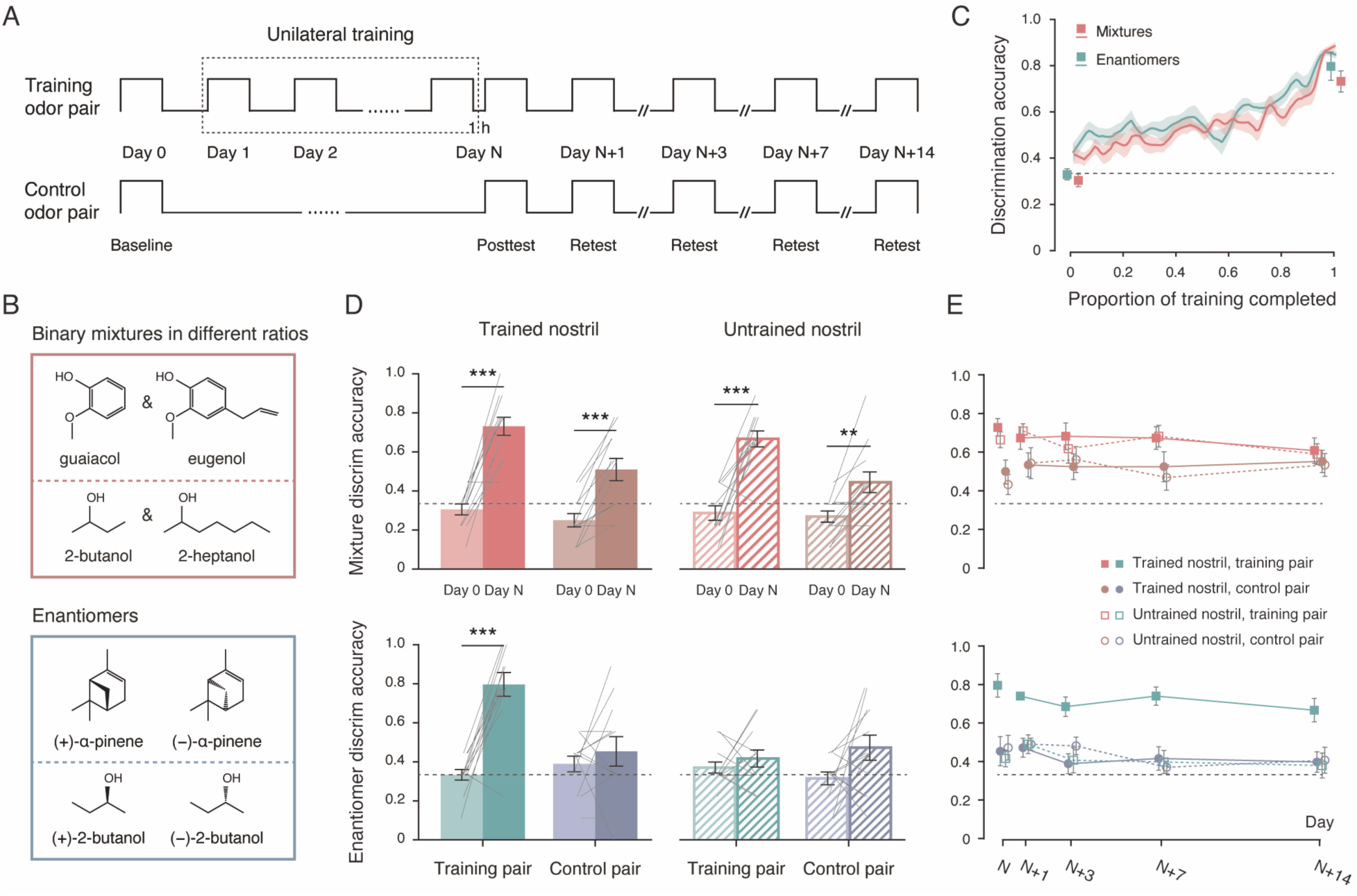
Experiment 1: distinct patterns of specificity and transfer in mixture discrimination learning and enantiomer discrimination learning. (A) Schematic illustration of the experimental procedure, which consisted of three phases: baseline (Day 0), unilateral olfactory training (Day 1 to Day N), and post-training testing (Day N, N+1, N+3, N+7, and N+14). Participants were divided into two groups: one trained and tested with binary odor mixtures (mixture group) and the other with odor enantiomers (enantiomer group). Each participant received training in a specific nostril (trained nostril) using a specific odor pair (training pair). (B) Chemical structures of the olfactory stimuli. Top: constituents of two pairs of binary odor mixtures; bottom: two pairs of odor enantiomers. (C) Improvements in mixture discrimination (red) and enantiomer discrimination (green) over the course of training. Data points were interpolated (gridded interpolation) and averaged across participants. Squares indicate the mean discrimination accuracies in each group for the training pair of odors presented to the trained nostril at baseline and in the post-training test on Day N. (D) Discrimination accuracies at baseline (Day 0, lighter bars) and on Day N’s post-training test (darker bars) for the training pair (red and green bars) and the control pair (brown and blue bars) presented to the trained (solid bars) and untrained (striped bars) nostrils in the mixture group (top) and the enantiomer group (bottom). Gray solid lines represent individual participants. (E) Discrimination accuracies in the post-training test sessions on Day N, N+1, N+3, N+7, and N+14 for the training pair and the control pair presented to the trained and untrained nostrils in the mixture group (top) and the enantiomer group (bottom). Black dashed lines: chance level (1/3). Shaded areas and error bars: SEMs. **: p < 0.01, ***: p ≤ 0.001.

The olfactory stimuli for the mixture group consisted of two pairs of binary mixtures (Figure 1B): a:b and b:a mixtures of guaiacol (1% v/v in propylene glycol) and eugenol (1% v/v in propylene glycol), and a:b and b:a mixtures of 2-butanol (1% v/v in propylene glycol) and 2-heptanol (1% v/v in propylene glycol). For three participants, the ratio a:b was 9:11, and for the remaining nine participants, it was 7:9. Each mixture comprised constituents similar in structure, comparable in olfactory intensity and valence (intensity: ts_23_ = 0.57 and 1.68, ps = 0.58 and 0.11 for guaiacol and eugenol and for 2-butanol and 2-heptanol, respectively; valence: ts_23_ = −0.84 and 0, ps = 0.41 and 1, respectively; ratings provided by an independent panel of 24 odor judges; Figure S1A), yet dissimilar in odor quality (mean similarity ratings ± SDs = 3.04 ± 1.85 and 2.67 ± 1.52, respectively, on a 7-point Likert scale where 7 marks extremely similar). Although the phenols guaiacol and eugenol are structurally distinct from the aliphatic alcohols 2-butanol and 2-heptanol, there was no overall difference in pairwise perceptual (dis)similarity between the structurally similar (either both phenols or both aliphatic alcohols, 2.85 ± 1.49) and dissimilar (one phenol, the other aliphatic alcohol, 2.86 ± 0.90) compounds (t_23_ = −0.035, p = 0.97; Figure S1B). Five of the participants used the phenolic mixtures as the training pair and the alcoholic mixtures as the control pair, with this reversed for the other seven. The olfactory stimuli for the enantiomer group consisted of two pairs of structurally unrelated odor enantiomers (Figure 1B) (Feng & Zhou, 2019): the enantiomers of α-pinene and those of 2-butanol. Each pair served as the training pair for half of the participants and as the control pair for the other half.

Throughout the experiment, olfactory discrimination was assessed unilaterally using a triangle task. In each trial, blindfolded participants sampled three bottles one at a time by sniffing with one nostril while keeping the other nostril pinched shut. Two of these bottles contained the same odorant and one contained a different odorant. They then identified which bottle smelled different from the other two (chance = 1/3). Feedback was provided on a trial-by-trial basis during the training phase (Roelfsema et al., 2010) but not during baseline (Day 0) or post-training testing. Training was conducted in the left nostril for roughly half of the participants in each group and in the right nostril for the other half, with one session per day on consecutive days (Day 1 to Day N, 12 trials per session). Only the training pair of mixtures or enantiomers was presented during these sessions. Training concluded when unilateral discrimination accuracy for the training pair exceeded 83% (≥ 10/12) on two consecutive days (Day N-1 and Day N). Post-training testing first took place 1 hour after the final training session (on Day N), followed by additional testing after 1, 3, 7, and 14 days (Day N+1, N+3, N+7, and N+14). Discriminations for both the training pair and the control pair were assessed for each nostril at baseline and post-training testing (9 trials per pair per nostril for a total of 36 trials per session).

At baseline, participants in both groups demonstrated chance-level performance (sums of correct discrimination = 120 and 152, out of 36 × 12 = 432 trials per group, binomial test ps = 0.99 and 0.22, for the mixture group and the enantiomer group, respectively), irrespective of the odor pairs’ identity (phenolic mixture vs. alcoholic mixture: F_1, 11_ = 1.92, p = 0.19; enantiomers of α-pinene vs. those of 2-butanol: F_1, 11_ = 0.88, p = 0.37) and whether the odors were presented to the left or right nostril (Fs_1, 11_ = 0.084 and 0.77, ps = 0.78 and 0.40, respectively). The number of sessions required to reach the training criterion ranged from 7 to 28 in both groups, with no difference between them (t_22_ = 1.32, p = 0.20). No sex difference was observed (t_22_ = 1.46, p = 0.16). The average learning curves for the two groups showed similar continuous improvements from the chance-level, indicating learning retention from one daily session to the next (Karni & Sagi, 1993) (Figure 1C). In other words, despite a large interindividual variation in the efficiency of olfactory perceptual learning, the discrimination of the mixtures and that of the enantiomers seemed comparable in task difficulty.

An omnibus ANOVA on the post-training discrimination accuracies on Day N, with role of odor pair (training vs. control) and nostril of presentation (trained vs. untrained) as the within-subjects factors and group (mixture vs. enantiomer) as the between-subjects factor, showed a robust three-way interaction between these factors (F_1, 22_ = 19.96, p < 0.001, η^2^_p_ = 0.48). Further comparisons of baseline and post-training discriminations within and between groups revealed comparable perceptual gains for the training pair presented to the trained nostril in both groups (t_22_ = −0.45, p = 0.66) –– the discrimination accuracies increased more than two-fold on average from Day 0 to Day N (from 0.32 to 0.76, t_23_ = 10.93, p = 1.40 × 10^-10^, Cohen’s d = 2.23) with no sex difference (t_22_ = 0.22, p = 0.83). However, the patterns of specificity/transfer diverged considerably between the mixture and enantiomer groups (Figure 1D).

In the mixture group, significant improvements in discrimination were also observed for the control pair presented to the trained nostril, the training pair presented to the untrained nostril, and the control pair presented to the untrained nostril (ts_11_ = 5.90, 6.29, and 3.51, ps < 0.001, < 0.001, and = 0.005, Cohen’s ds = 1.70, 1.82, and 1.01, respectively). The perceptual gains were less pronounced for the control pair relative to the training pair (increments in discrimination accuracy: 0.22 vs. 0.40; main effect of odor pair: F_1, 11_ = 25.88, p < 0.001, η^2^_p_ = 0.70), but did not differ between the trained and untrained nostrils (Fs_1, 11_ = 1.77 and 0.20, ps = 0.21 and 0.66, for the main effect of nostril and interaction between odor pair and nostril, respectively). Overall, the participants became significantly above chance in discriminating not only the training pair but also the control pair of odor mixtures (mean accuracies = 0.70 and 0.48 vs. chance = 0.33, ts_11_ = 9.69 and 3.22, ps < 0.001 and = 0.008, Cohen’s ds = 2.80 and 0.93, respectively) independent of the nostril of presentation (ts_11_ = 1.47 and 1.00, ps = 0.17 and 0.34, respectively). These results indicated that learning of mixture discrimination completely transferred across nostrils. Critically, it effectively generalized to untrained mixtures that were unrelated to the training material in structure and odor quality.

For the enantiomer group, learning was restricted to the training pair presented to the trained nostril. No significant perceptual gain was observed in any other condition (ts_11_ = 0.80, 0.83, and 1.96, ps = 0.44, 0.42, and 0.076, for control pair presented to the trained nostril, training pair presented to the untrained nostril, and control pair presented to the untrained nostril, respectively), consistent with earlier findings on chiral discrimination learning (Feng & Zhou, 2019).

### Persistence of the learning effects

To assess the persistence of the learning effects described above, we conducted separate repeated-measures ANOVAs on the post-training discrimination accuracies of each group, using role of odor pair (training vs. control), nostril of presentation (trained vs. untrained), and test session (Day N, N+1, N+3, N+7, and N+14) as the within-subjects factors. In both groups, we found no significant main effect of test session (Fs_4, 44_ = 0.33 and 2.48, ps = 0.86 and 0.058, for the mixture and enantiomer groups, respectively) or any interaction between test session and the other factors (ps > 0.1). In the mixture group, a significant main effect of odor pair was obtained (F_1, 11_ = 14.95, p = 0.003, η^2^_p_ = 0.58), with no main effect of nostril or interaction between odor pair and nostril (Fs_1, 11_ = 0.53 and 0.001, ps = 0.48 and 0.97, respectively). However, in the enantiomer group, there was a significant interaction between odor pair and nostril (F_1, 11_ = 22.92, p < 0.001, η^2^_p_ = 0.68), in addition to significant main effects of odor pair and nostril (Fs_1, 11_ = 19.18 and 32.81, ps = 0.001 and < 0.001, η^2^_p_s = 0.64 and 0.75, respectively). Thus, discrimination performances were stable in both groups and their respective patterns of specificity/transfer were retained throughout all test sessions (Figure 1E). Two weeks post-training (Day N+14), participants in the mixture group continued to discriminate not only the training pair but also the control pair of odor mixtures significantly above chance (ts_11_ = 5.26 and 4.37, ps < 0.001 and = 0.001, Cohen’s ds = 1.52 and 1.26, respectively), regardless of the nostril of presentation (ts_11_ = 0.29 and 0.28, ps = 0.78 and 0.78, respectively). Conversely, those in the enantiomer group still showed supra-chance discrimination only for the training pair of enantiomers presented to the trained nostril (ts_11_ = 5.45, p < 0.001, Cohen’s d = 1.57; ps > 0.25 for all other conditions).

Aside from effects on discrimination, prolonged unilateral olfactory training led to odor- and nostril-specific adaptation in both groups. At baseline, the participants perceived the training pair and the control pair of odorants presented to each nostril as similarly intense (Fs_3, 33_ = 0.30 and 0.13, ps = 0.83 and 0.94, for the mixture and enantiomer groups, respectively). However, at the post-training test on Day N, they rated the training pair presented to the trained nostril as significantly weaker than the training pair presented to the untrained nostril, the control pair presented to the trained nostril, as well as the control pair presented to the untrained nostril (ps < 0.01, Figure S2A). There was no difference between the latter three conditions (Fs_2, 22_ = 0.43 and 0.048, ps = 0.66 and 0.95, respectively). This adaptation effect gradually diminished over time (F_5.18, 62.16_ = 3.81, p = 0.004, η^2^_p_ = 0.24, Figure S2B). Two weeks post-training (Day N+14), perceived odor intensities were no longer affected by the nostril of presentation (ts_15_ = −0.55 and 1.38, ps = 0.59 and 0.19 for the training and control pairs, respectively), while learned abilities of odor discrimination were retained (Figure 1E).

### Verification of long-term transfer of olfactory learning with new odor mixtures

It could be argued that the transfer of odor discrimination learning between the phenolic mixtures and alcoholic mixtures, as observed in Experiment 1, was facilitated by a common element between the two –– the hydroxyl group. To investigate whether learning transfer occurs between odor mixtures that do not share a common functional group, we introduced two new pairs of odor mixtures in Experiment 2 (Figure 2A). One pair consisted of a:b and b:a mixtures of acetophenone (1% v/v in propylene glycol) and 2-octanone (1% v/v in propylene glycol), and the other of a:b and b:a mixtures of methyl salicylate (1% v/v in propylene glycol) and isoamyl butyrate (1% v/v in propylene glycol). The constituents of each binary mixture, one aromatic and the other aliphatic, were selected to be structurally more distinct compared to those in Experiment 1. Based on ratings from a new panel of 24 odor judges (Figure S3A), acetophenone and 2-octanone were comparable in odor intensity and valence (ts_23_ = −1.56 and 0.30, ps = 0.13 and 0.76, respectively), whereas methyl salicylate was equally intense to but less pleasant than isoamyl butyrate (ts_23_ = −1.07 and −2.64, ps = 0.29 and 0.015, respectively). All compounds were dissimilar in terms of odor quality (mean pairwise similarity ratings < 3.17 for any two compounds), with no difference in pairwise perceptual (dis)similarity between those within the same functional group (either both ketones or both esters, 2.54 ± 1.27) and those in different functional groups (one ketone, the other ester, 2.45 ± 0.83) (t_23_ = 0.45, p = 0.65; Figure S3B). Using the same procedures as in Experiment 1 (Figure 1A), we trained and tested another 12 participants for odor mixture discrimination unilaterally. The mixing ratio a:b was 9:11 for 9 participants and 7:9 for 3 participants.

**Figure 2.**
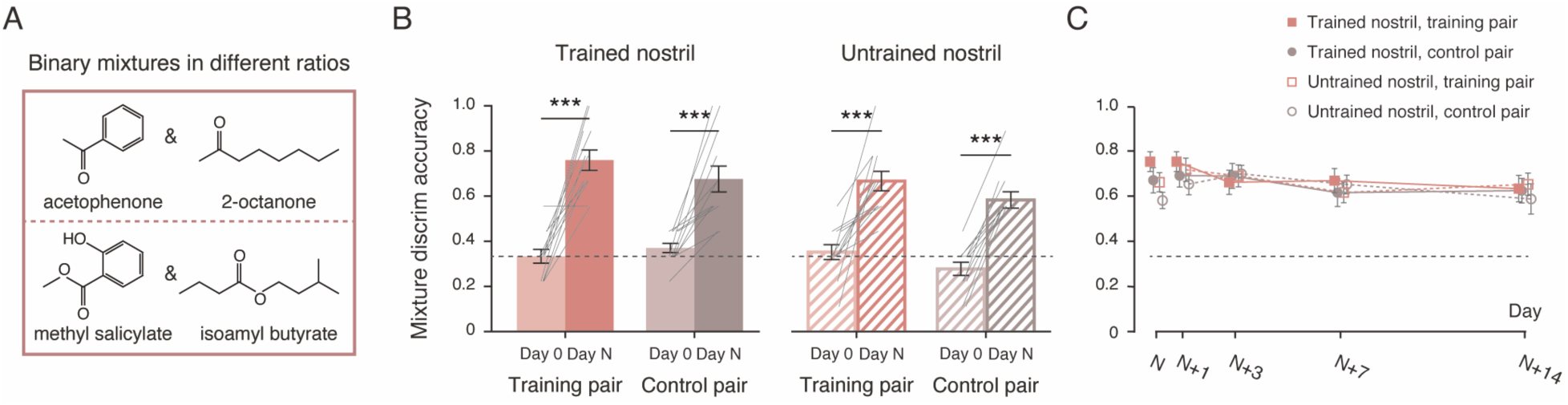
Experiment 2: replication of long-term transfer of mixture discrimination learning. (A) Chemical structures of the constituents of two new pairs of binary odor mixtures used in Experiment 2. (B) Discrimination accuracies at baseline (Day 0, lighter bars) and on Day N’s post-training test (darker bars) for the training pair (red bars) and the control pair (brown bars) of odor mixtures presented to the trained (solid bars) and untrained (striped bars) nostrils. Gray solid lines represent individual participants. (C) Discrimination accuracies in the post-training test sessions on Day N, N+1, N+3, N+7, and N+14 for the training pair and the control pair presented to the trained and untrained nostrils. Black dashed lines: chance level (1/3). Error bars: SEMs. ***: p ≤ 0.001.

At baseline, the participants performed at chance level in discriminating the binary odor mixtures (sum of correct discrimination = 144 out of 36 × 12 = 432 trials, binomial test p = 0.52), regardless of the type of mixtures presented (ketones vs. esters: F_1, 11_ = 0, p > 0.99) or whether the odors were presented to the left or right nostril (F_1, 11_ = 0.077, p = 0.79). Subsequently, they were randomly assigned to receive training in either the left nostril (5 participants) or the right nostril (7 participants), with either the ketone mixtures (7 participants, with the ester mixtures serving as the control pair) or the ester mixtures (5 participants, with the ketone mixtures as the control pair). It took them between 16 and 34 sessions (N = 16 to N = 34) to reach the training criterion, with no difference between the two sexes (t_10_ = 0.45, p = 0.67).

Comparisons of the performances at baseline and the post-training test on Day N showed substantially improved discriminations not only for the training pair presented to the trained nostril (t_11_ = 7.37, p < 0.001, Cohen’s d = 2.13), but also for the control pair presented to the trained nostril, the training pair presented to the untrained nostril, and the control pair presented to the untrained nostril (ts_11_ = 5.25, 5.17, and 7.84, ps < 0.001, Cohen’s ds = 1.51, 1.49, and 2.26, respectively; Figure 2B) in both men and women (F_1, 10_ = 0.094, p = 0.77). Specifically, at the post-training test on Day N, individuals trained with the ketone mixtures demonstrated above-chance discriminations for both the ketone mixtures and the ester mixtures presented to both the trained and untrained nostrils (mean accuracies = 0.75, 0.71, 0.65, and 0.52 vs. chance = 0.33,

Wilcoxon Zs = 2.37, 2.38, 2.38, and 2.53, ps = 0.018, 0.017, 0.017, and 0.011, respectively); those trained with the ester mixtures also demonstrated above-chance discriminations for both the ester mixtures and the ketone mixtures presented to both the trained and untrained nostrils (mean accuracies = 0.78, 0.60, 0.71, and 0.67 vs. chance = 0.33, Wilcoxon Zs = 2.06, 2.03, 2.02, and 2.04, ps = 0.039, 0.042, 0.043, and 0.041, respectively). Thus, the learning effect significantly transferred across nostrils and between the ketone mixtures and ester mixtures that were perceptually dissimilar and shared no common functional group, echoing the results obtained with the phenolic and alcoholic mixtures in Experiment 1. Furthermore, the discrimination accuracies for the untrained mixture pair observed here exceeded those in Experiment 1 by 15% (0.63 vs. 0.48, t_22_ = 2.55, p = 0.018, Cohen’s d = 1.04), suggesting that structural diversity of the training material could enhance generalization of the learning effect. In parallel, training also resulted in a specific adaptation to the training pair of mixtures presented to the trained nostril (Figure S4A), which was perceived as significantly less intense than the training pair presented to the untrained nostril, the control pair presented to the trained nostril, and the control pair presented to the untrained nostril at the post-training test on Day N (ps < 0.05), but not at baseline (F_3, 33_ = 0.60, p = 0.62).

Similar to Experiment 1, discrimination performances, along with the pattern of transfer, remained stable over the 5 post-training test sessions spanning two weeks (Day N, N+1, N+3, N+7, and N+14, F_4, 44_ = 1.42, p = 0.24, interactions with other factors: ps > 0.1; Figure 2C). Overall, discrimination accuracies were numerically lower for the control pair compared to the training pair of odor mixtures, but this difference was not statistically significant (F_1, 11_ = 3.50, p = 0.088), reflecting substantial generalization of the learning effect. These accuracies were comparable between the trained and untrained nostrils (Fs_1, 11_ = 2.16 and 0.019, ps = 0.17 and 0.89 for the main effect of nostril and interaction between nostril and odor pair, respectively), indicating a complete learning transfer across nostrils. Meanwhile, training-induced olfactory adaptation gradually lessened (F_12, 132_ = 2.75, p = 0.002, η^2^_p_ = 0.20, Figure S4B). Two weeks post-training (Day N+14), participants continued to discriminate not only the training pair but also the control pair of odor mixtures significantly above chance (ts_11_ = 6.30 and 4.93, ps < 0.001, Cohen’s ds = 1.82 and 1.42, respectively), irrespective of the nostril of presentation (ts_11_ = −0.39 and 0.74, ps = 0.70 and 0.47, respectively). Their perceptions of odor intensity were no longer affected by the nostril of presentation or whether the mixtures had been used for training (main effects: Fs_1, 11_ = 0.60 and 0.026, ps = 0.46 and 0.87, respectively, interaction: F_1, 11_ = 0.061, p = 0.81).

## Discussion

Generalization enables the transfer of learning from specific parameters to broader, new situations. This typically involves the transfer of abstract knowledge or patterns of characteristics from specific circumstances (Kellman & Massey, 2013; Tenenbaum et al., 2011). In this study, we employed the same protocol but introduced different types of odors for unilateral olfactory discrimination training. We observed contrasting patterns of specificity and transfer. Learning was confined to the training pair presented to the trained nostril in individuals trained with odor enantiomers. However, it completely transferred to the untrained nostril and effectively (though not entirely) generalized to other structurally and perceptually unrelated mixtures in those trained with odor mixtures. The structural diversity within the mixtures used for training seemed to further enhance the generalization of the learning effect. These results were obtained despite the highly similar subjective experiences (to discriminate between two highly similar odors), task difficulties, and learning curves between participants trained with odor mixtures and those with enantiomers. Moreover, the learning effects were long-lasting, persisting for at least two weeks in both cases, and were not dependent on olfactory adaptation or participants’ sex.

Anatomically, olfactory inputs to the two nostrils remain largely separated up to the primary olfactory cortex, which includes the anterior olfactory nucleus, olfactory tubercle, piriform cortex, several amygdala subnuclei, and rostral entorhinal cortex (Gottfried & Zald, 2005). The piriform cortex is known to transform the chemical feature space inherited from the olfactory bulb into an odor quality space characterized by coding sparseness (Bekkers & Suzuki, 2013; Howard et al., 2009; Pashkovski et al., 2020; Stettler & Axel, 2009). Downstream relays from the primary olfactory cortex to the orbitofrontal cortex, lateral and basolateral amygdala, hippocampus, as well as a range of other structures in the limbic and paralimbic cortices link olfactory inputs to systems associated with affect, learning, memory, and value appraisal (Gottfried, 2010; Gottfried & Zald, 2005). The nostril- and structure-specificity of enantiomer discrimination learning likely reflects plasticity in or upstream of the olfactory bulb (Feng & Zhou, 2019). Conversely, the internarial transfer of learning observed with mixture discrimination suggests that learning occurs at a stage where convergence of binaral input has occurred –– i.e., at a stage higher than the primary olfactory cortex. The generalization of this learning to structure- and odor quality-wise unrelated mixtures further indicates the involvement of abstract representations of discrimination boundaries for inputs varying in odor quality. A potential locus of plasticity underlying such generalization is the orbitofrontal cortex, which is the main neocortical target of primary olfactory areas (Gottfried, 2010). It has been suggested that this region integrates olfactory perceptual evidence toward a criterion bound for discrimination (Bowman et al., 2012). Presumably, prolonged training with a pair of odor mixtures refines the representation of perceptual evidence (i.e., relative strengths of odor notes) in the orbitofrontal cortex and modifies the discrimination boundary (Qamar et al., 2013). The application of this optimized boundary subsequently produces a general improvement in odor mixture discrimination that extends beyond the training pair of mixtures and the trained nostril.

The differences in specificity and transfer between learning to discriminate mixtures and enantiomers cannot be attributed to a generic effect of attention, adaptation, reinforcement, task difficulty, or the extent of training (Ahissar & Hochstein, 1997, 2004; Harris et al., 2012; Karni & Sagi, 1993; Roelfsema et al., 2010). Instead, they align with the conceptual framework of the reweighting model of perceptual learning (Lu & Dosher, 2022) and the notion that learning relies on a distribution of plasticity across the brain (Maniglia & Seitz, 2018). It is plausible that the learned weight structure for discriminating between two odor mixtures (i.e., relative strengths of odor notes) roughly matches the desired weight structure for discriminating other mixtures, thus allowing for transfer. Conversely, the learned weight structure in the case of enantiomer discrimination (i.e., configuration of the chiral center) is incompatible with that required for discriminating other structurally-dissimilar enantiomers, leading to high specificity. In simpler terms, learning to discriminate different odors spontaneously engages plasticity at distinct stages of olfactory processing.

Our findings bear significant practical implications. They demonstrate the generalizability (with mixture discrimination) and persistence of learning-induced olfactory gain, thereby supporting the potential value of olfactory training as a therapeutic intervention for olfactory dysfunction and as a method to acquire olfactory expertise. Importantly, they call for the utilization of more complex odors and tasks (as opposed to single compounds and passive exposure) to facilitate the generalization of learning effects beyond the training setting. Given the unique anatomical proximity between secondary olfactory areas and regions implicated in emotion and memory (Gottfried, 2010; Gottfried & Zald, 2005), our results also raise the intriguing possibility that olfactory training, when conducted with carefully designed material and protocol, may act on emotional and mnemonic functions. This will be subject to future studies.

## Materials and methods

### Participants

A total of 36 healthy non-smokers completed the unilateral training of odor discrimination, which lasted for an average of 20 days: 24 (12 females, 22.3 ± 1.4 years) in Experiment 1 and 12 (6 females, 25.1 ± 1.5 years) in Experiment 2. The sample size of each group (n = 12) was comparable to those in previous perceptual learning studies (Dalton et al., 2002; Feng & Zhou, 2019; Mainland et al., 2002). An additional 48 healthy non-smokers provided odor ratings for the constituents of mixtures used in Experiment 1 (n = 24, 12 females, 25.3 ± 2.3 years) and Experiment 2 (n = 24, 12 females, 25.2 ± 2.2 years). All participants reported having a normal sense of smell and no respiratory allergies or upper respiratory infections at the time of testing. They provided written informed consent to participate in procedures approved by the Institutional Review Board at the Institute of Psychology, Chinese Academy of Sciences (H18029).

### Olfactory stimuli

In Experiment 1, the olfactory stimuli consisted of two pairs of binary mixtures and two pairs of enantiomers (Figure 1B): a:b and b:a mixtures of guaiacol (CAS number: 90-05-1, 1% v/v) and eugenol (97-53-0, 1% v/v), a:b and b:a mixtures of 2-butanol (78-92-2, 1% v/v) and 2-heptanol (543-49-7, 1% v/v), (+)-α-pinene (7785-70-8, 1% v/v) and (-)-α-pinene (7785-26-4, 1% v/v), as well as (+)-2-butanol (4221-99-2, 1% v/v) and (-)-2-butanol (14898-79-4, 1% v/v). In Experiment 2, the olfactory stimuli comprised two new pairs of binary mixtures (Figure 2A): a:b and b:a mixtures of acetophenone (98-86-2, 1% v/v) and 2-octanone (111-13-7, 1% v/v), and a:b and b:a mixtures of methyl salicylate (119-36-8, 1% v/v) and isoamyl butyrate (106-27-4, 1% v/v). All compounds were dissolved in propylene glycol. The mixing ratio a:b was 9:11 for 3 participants in Experiment 1 and 9 participants in Experiment 2, and 7:9 for 9 participants in Experiment 1 and 3 participants in Experiment 2. The constituents of the mixtures, namely guaiacol, eugenol, 2-butanol, and 2-heptanol in Experiment 1, and acetophenone, 2-octanone, methyl salicylate, and isoamyl butyrate in Experiment 2, were evaluated for their odor characteristics prior to the main experiments (see Supplemental Methods).

The olfactory stimuli were presented in identical 40 ml polypropylene jars. Each jar contained 10 ml of clear liquid and was fitted with either one (for unilateral training and testing) or two (for odor evaluations) Teflon nosepieces. Participants were instructed to inhale through the nosepieces and exhale through the mouth. Fresh odor solutions were prepared every other day during the period of data collection.

### Procedure

We adopted procedures similar to those outlined in a previous study on olfactory perceptual learning of chiral discrimination (Feng & Zhou, 2019). Specifically, Experiment 1 was composed of three phases: baseline, unilateral training, and post-training testing (Figure 1A). At baseline (Day 0), participants, blindfolded, completed 8 trials of a unilateral odor intensity rating task, followed by 36 trials of a unilateral odor discrimination task. In each trial, they were instructed to pinch one nostril shut and not release it until the completion of that trial. Odors were presented to the open nostril by the experimenter. For half of the participants, the presented odors were two pairs of binary odor mixtures: guaiacol and eugenol, and 2-butanol and 2-heptanol, both in the ratios of a:b and b:a. For the other half, the presented odors were the enantiomers of ɑ-pinene and those of 2-butanol. In the odor intensity rating task, participants were presented with one odor per trial. They rated its intensity on a 7-point Likert scale, with 7 representing “extremely intense”. The two odors within a pair were presented in random order in 4 consecutive trials with the nostril of presentation alternated between trials (1 trial per odor per nostril). In the odor discrimination task, participants were presented with three odors –– two identical and one different –– in each trial, one after another in random order. They were asked to identify the one that smelled different from the other two. Each pair was presented in 18 consecutive trials with the nostril of presentation alternated between trials (9 trials per pair per nostril). The order of odor pairs was counterbalanced across participants. There was a 30-s break between two trials and a 10-min break after the first 18 trials of the odor discrimination task. No feedback was provided. Participants with discrimination accuracies ≤ 5/9 (vs. chance = 1/3) for each odor pair were invited for olfactory training.

Training started the day after the baseline session (Day 1) and was conducted at approximately the same time each day on consecutive days (Day 1 to Day N). Each training session comprised 12 trials of the unilateral odor discrimination task conducted in a specific nostril (trained nostril) using a specific odor pair (training pair), with a 30-s break between two trials. Immediate feedback was provided after each trial. For the mixture group, 5 participants were trained with the phenolic mixtures and 7 were trained with the alcoholic mixtures. For the enantiomer group, 6 participants were trained with the enantiomers of ɑ-pinene and 6 were trained with the enantiomers of 2-butanol. Training was conducted in the left nostril for 7 participants in the mixture group and 6 participants in the enantiomer group, and in the right nostril for the remaining participants. It concluded when the discrimination accuracy reached 10/12 or higher on two consecutive days (Day N-1 and Day N).

The first post-training testing session took place one hour after the final training session on Day N, using materials and procedures identical to those in the baseline session. Additional retests were carried out after 1, 3, 7, and 14 days (Day N+1, N+3, N+7, and N+14). The retest sessions were the same as the baseline session and the post-training test session on Day N except that not all participants performed the unilateral odor intensity rating task. On each of Day N+1, N+3, and N+7, 9 participants in the mixture group and 4 in the enantiomer group provided odor intensity ratings. On Day N+14, 12 participants in the mixture group and 4 in the enantiomer group provided intensity ratings.

Except for the olfactory stimuli used, the procedures in Experiment 2 were identical to those for the mixture group in Experiment 1. Specifically, the ketone mixtures served as the training pair and the ester mixtures served as the control pair for 7 participants. This was reversed for the remaining 5 participants. Training was conducted in the left nostril for 5 participants and in the right nostril for the other 7 participants. All participants completed the unilateral odor intensity rating task and the unilateral odor discrimination task at baseline (Day 0) and at the post-training test and retest sessions held on Day N, N+1, N+3, N+7, and N+14.

### Statistical analysis

Participants’ overall baseline performances in odor discrimination were compared against chance using binomial tests. Unilateral discrimination accuracies for various odor pairs, both at baseline and post-training tests, were analyzed using repeated-measures ANOVAs. Specifically, the ANOVAs examined: (1) the impact of odor pair identity (mixture groups: phenolic mixtures vs. alcoholic mixtures or ketone mixtures vs. ester mixtures, enantiomer group: enantiomers of α-pinene vs. those of 2-butanol) and side of presentation (left vs. right) on baseline discrimination accuracies in each group; (2) the influence of group (mixture vs. enantiomer), assigned role of odor pair (training vs. control), and nostril of presentation (trained vs. untrained) on post-training discrimination accuracies on Day N in Experiment 1; (3) the effect of post-training test session (Day N, N+1, N+3, N+7, and N+14), role of odor pair (training vs. control), and nostril of presentation (trained vs. untrained) on learning effects (i.e., post-training discrimination accuracies) in each group. In addition, a series of t-tests and Wilcoxon signed rank tests (non-parametric, suitable for small sample sizes) were conducted to compare discrimination accuracies or training-induced perceptual gains (calculated as the difference in discrimination accuracy between the post-training test on Day N and the baseline) between conditions and to compare discrimination accuracies against the chance-level (1/3). Unilateral odor intensity ratings for different odor pairs were similarly analyzed using repeated-measures ANOVAs and t-tests. In Experiment 1, both groups of participants showed nostril-specific adaptation to the training pair of odors at the post-training test on Day N. We subsequently combined the intensity rating data from the two groups on Day N, N+1, N+3, N+7, and N+14 to examine the change in this adaptation effect over time.

The effect sizes of ANOVAs and t tests were estimated using partial eta squared (η^2^_p_) and Cohen’s d, respectively. All t-tests were two-tailed.

## Acknowledgements

This work was supported by STI2030-Major Projects 2021ZD0204200 (W.Z.), the Chinese Academy of Sciences Grant JCTD-2021-06 (W.Z.), and the National Natural Science Foundation of China Grant 32430043 (W.Z.).

## Data availability

All data generated or analyzed during this study are included in the manuscript and supporting files; source data files have been provided for Figures 1 and 2.

